# Innate triggering and antiviral effector functions of activin A

**DOI:** 10.1101/2021.03.23.436626

**Authors:** K. Al-Hourani, N Ramamurthy, E Marchi, RM Eichinger, LN Lee, P Fabris, P Klenerman, H. Drakesmith

## Abstract

First-line defence against viral infection is contingent upon rapid detection of conserved viral structural and genomic motifs by germline-encoded pattern recognition receptors, followed by activation of the type I IFN system and establishment of an intracellular antiviral state. Novel antiviral functions of bone morphogenetic protein and related activin cytokines, acting in conjunction with, and independently of, type I IFN, have recently been described. Activin A mediates multiple innate and adaptive immune functions – including antiviral effects. However, how such effects are mediated and how activin might be triggered by viral infection have not been defined. Here we addressed this in vivo and in vitro, in humans and mice.

Transcriptomic analyses delineated strikingly congruent patterns of gene regulation in hepatocytes stimulated with recombinant activin A and IFNα *in vitro*. Activin A mRNA, encoded by *INHBA*, is induced upon activation of RIG-I, MDA5 and TLR7/8 viral nucleic acid sensors *in vitro*, across multiple cell lines and in human peripheral blood mononuclear cells. *In vivo*, infection of mice with influenza A also upregulated *Inhba* mRNA in the lung; this local upregulation of *Inhba* is retained in MAVS knockout mice, indicating a role for non-RIG-I-like receptors in its induction. Activin induction and signalling were also detectable in patients with chronic viral hepatitis.

Together, these data suggest Activin A is triggered in parallel with type I IFN responses and can trigger related antiviral effector functions. This model has implications for the development of targeted antiviral therapies, in addition to revealing novel facets of activin biology.

## Introduction

First identified as a regulator of ovarian folliculogenesis, activin A(1), a heterodimeric assembly of inhibin β_A_ subunits (encoded by the *INHBA* gene), is implicated in diverse biological processes. In terms of innate immune functionality, activin A is induced in peripheral blood mononuclear cells exposed to pro-inflammatory stimuli including TNFα, GM-CSF and IFNγ(2); neutrophil granules are also rich sources of pre-formed activin A protein(3). *In vivo*, the induction kinetics of activin A in response to systemic inflammatory stimuli have been examined following intravenous LPS exposure: serum activin A protein increases rapidly during endotoxaemic shock in mice, with administration of its soluble antagonist follistatin (FST) sufficient to reduce subsequent mortality(4). Clinically, in H1N1 influenza-infected patients admitted to intensive care, elevated serum levels of both activin A and FST were detected, and correlated with degree of respiratory distress(5).

Recent observations have shown an antiviral function of the TGFβ-superfamily cytokines the bone morphogenetic proteins (BMP) and the activins, both transduced via SMAD transcription factors, a phenomenon elucidated via examination of the mutually antagonistic interactions between Hepatitis C Virus (HCV) and the BMP/SMAD signalling axis. Activin A exerts dose-dependent antiviral effects against an *in vitro* HCV genomic replicon(6). Additionally, *in vitro* antiviral functions of activin A against Zika virus, a flavivirus akin to HCV, and also against Hepatitis B Virus (HBV), a structurally distinct hepadnavirus(6) have been reported.

While activin proteins can mediate antiviral effects against multiple viruses *in vitro*, upregulation of activins in response to viral infection, akin to the rapid induction of type I IFN, has not previously been described. Three classes of pattern recognition receptor (PRR) sense viral nucleic acids: RIG-I-like receptors (RLR); Toll-like receptors (TLR); and the cGAS-STING axis for detection of cytosolic DNA. RIG-I, the prototypic RLR detects 5’-triphosphate and diphosphate moieties associated with non-self RNA(7,8). The RLR family also includes MDA5, a RIG-I paralogue demonstrating length-dependent activation by dsRNA(9).RLR activation drives the oligomerization of the adaptor protein MAVS, the essential factor for downstream type I IFN induction(10).

In humans, four endosomal TLRs are sensitive to non-self nucleic acids:TLR3, TLR7, TLR8 and TLR9(11). TLRs comprise an ectodomain conferring ligand specificity;a transmembrane region; and a cytosolic Toll/IL-1 receptor that ultimately activates IRAK, IKK and TBK1 kinases (11).

In this study, we first used microarray analysis and RNA sequencing to analyse the effects of activin A upon hepatocytes at the transcriptional level and examine its intersection with the type I IFN axis. We next addressed whether activin A transcription is induced upon both PRR activation and viral infection, both *in vitro* and *in vivo*, in addition to in part delineating the mechanistic basis for this phenomenon. Overall these data indicate that Activin plays a role in virus infections as part of the innate response.

## Results

### Activin A stimulation upregulates immune pathways in HepaRG cells

As shown in **Fig 1A**, in order to better understand the effect of Activin A on hepatocytes, we used microarray analysis of the RNA isolated from HepaRG cells activated with 10nM Activin A for 24 hrs. Differential analysis of the expression data show that 184 genes were upregulated and 168 number of genes were down-regulated with a minimum of 1 log fold change. **Table 1** shows the top 40 list of differentially regulated genes (limma, paired t test p<0.001 lfc>1). Metacore analysis (Clarivate Analytics) of the differentially regulated genes show that genes in Type 1 alpha/beta signalling pathway were predominantly upregulated, followed by MHC Class I presentation and Response to RNA viral infection **(Fig 1 B)**. However, Activin A seems to be downregulating genes in the cell cycle pathway **(Fig 1C)**.

**Figure 1.**
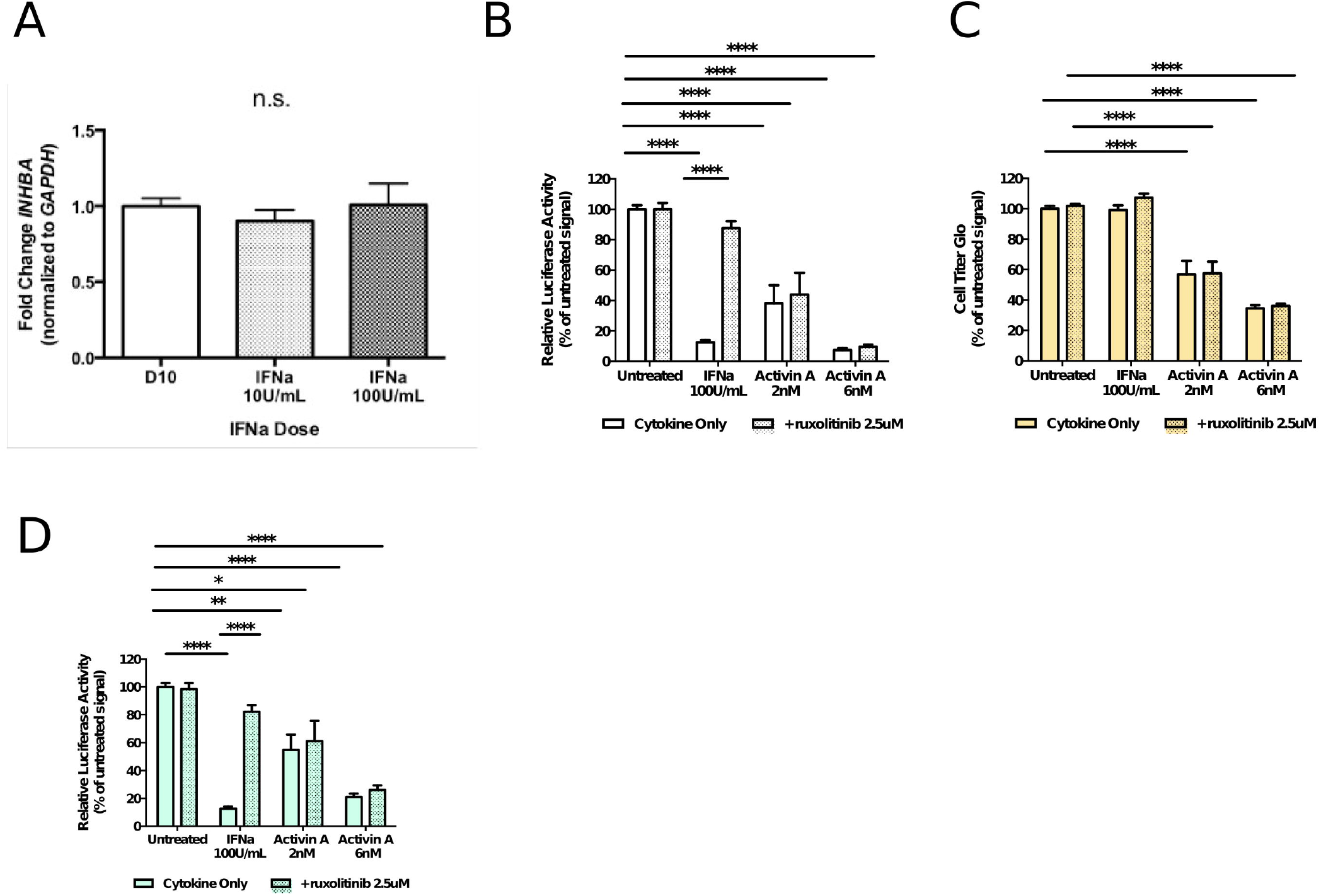
**A)** Diagram representing experimental cell culture procedure on stimulation of HepaRG cells with Activin A. **B)C)** Metacore analaysis performed on differentially regulated genes identified by microrray analysis of RNA from HepaRG cells stimulated with 10nM activin A over 24 hours. Figure 1B highlights pathways mapped to the 184 signficantly upregulated genes; figure 1C highlights those mapped to the 168 significantly downregulated transcripts. **D)** Time course over a period of 72 hours of HepaRG cells stimulated with Activin A (10nM), with relative gene expression levels of various ISGs shown compared to unstimulated cells. n=3 independent experiments conducted in triplicate. **E)** Comparative analysis of gene expression in different hepatocyte cell lines (HuH7, HepaRG, HHL12) at 24 hours following stimulation with Activin A (10nM). n=3 independent experiment conducted in triplicate.

### Activin A stimulates ISGs in hepatocyte cell lines

Intrigued by the ability of Activin A to up-regulate ISG in the absence of Interferons, we looked to address whether Activin A increases ISG levels by regulating IFN expression in the HepaRG hepatocyte cell line over a 72 hr period (**Fig. 1D**). qPCR analysis validated the increase in ISG expression observed in microarray and RNA sequencing. HepaRG cells stimulated with Activin A showed an increase in expression of various ISGs relative to un-stimulated cells **(Fig 1D)**. However, when tested on other cell lines, namely Huh7 hepatoma-derived cells and the immortalised transformed hepatocyte cell line HHL12 (12), we observed that the pattern on gene expression over time is not consistent and individual cell lines behave differently to Activin A in terms of ISG expression **(Fig. 1E)** The primer pairs used for the qPCR are detailed in the supplementary information **(Supplementary Table 1)**.

### RNA-sequencing confirms results of microarray data analysis on hepatocytes stimulated with Activin A

We corroborated our data obtained by microarray analysis by performing a new RNA sequencing experiment on a separate experiment with hepatocytes stimulated with Activin A. We then performed a Gene set enrichment analysis (GSEA) comparing both experiments. GSEA shows high correlation in gene expression by both methods of analysis. The entire set of Activin vs Untreated up-regulated genes detected in the RNA-Seq experiment (n=2571, data processed with DESeq2 package(13)) were found highly enriched compared to the microarray data (Fig 2a).

**Figure 2:**
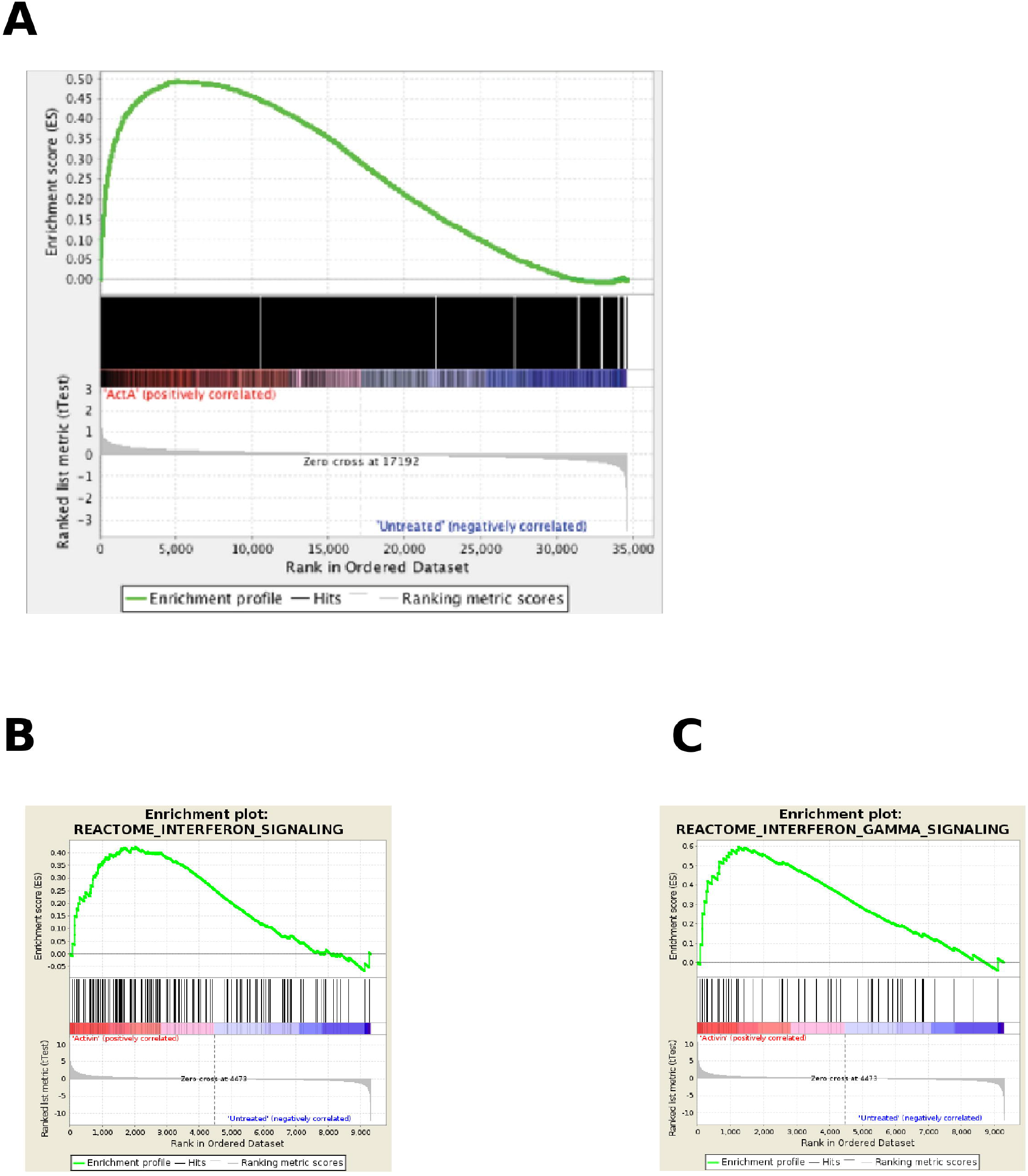
**A)** Gene Set Enrichment Analysis (GSEA) showing consistency of results between microrray and RNA-sequencing experiments interrogating transcriptomes elicited by treatment of HepaRG hepatoma-derived cells with activin A 10nM over a period of 24 hours. The whole set of up-regulated genes (log FC >0,p value > 0.1, n=2571) between Activin vs Untreated in RNA-seq were found significantly enriched (FDR = 0.04, p << 0.01) in Activin-treated samples derived from the equivalent microarray experiment described in Figure 1A (Activin vs Untreated condition). **B)C)** GSEA showing enrichment of Interferon gene pathways in transcriptomics from Activin treated samples. Based on 110 Interferon-related signatures selected from MsigDb, two representative Interferon Gene Sets were defined, corresponding to Interferon Signalling (Figure 2B) and Interferon Gamma Signalling. Both of these gene sets were founds to be significantly enriched in RNA-derived from HepaRG cells stimulated with activin A 10nM over 24 hours (Figure 2A: FDR = 0.19, p=0.026; Figure 2B: FDR = 0.019, p=0.004 respectively.

To show that ISG were upregulated in response to activin induction, we performed a GSEA using a large set of Interferon signatures publicly available in MsigDB database(14). We selected 110 gene sets querying all curated databases (e.g Reactome, GO, etc.) and reported gene expression data from immunologic and cancer studies (e.g. Up-regulated genes on cells stimulated with IFNa) in MsigDB. We found that 88 of all 110 gene sets were up-regulated in Activin treated samples, of which 24 gene sets were significantly enriched at FDR < 25%, 10 gene sets were at nominal p-value < 0.01, and 19 gene sets were significant at nominal p-value < 0.05. The remaining 22/110 gene sets were enriched in Untreated samples without any statistical significance. **Fig 2b** and **Fig 2c** shows two examples of public IFN signatures enriched in up-regulated genes in our Activin vs Untreated samples (in RNA-Seq data after *voom* transformation in *limma* R package(15)).

### Activity of Activin A is independent of IFNα

We have previously shown that recombinant Activin A protein exerts a dose dependent antiviral effect(6). We first explored the transcriptional pathways and show that Activin A is able to induce ISG signalling in the presence of B18R, a type I IFN binding protein derived from Western Reserve Strain Vaccinia virus, in Huh7 cells (**Fig. 3A**). However, B18R is able to inhibit RNA expression of ISG in response to IFNα (**Fig. 3B**). In addition, supernatants of hepatocyte (Huh7) cells stimulated with Activin A did not show the presence of IFNA2 (Fig 3C) supporting the view that Activin A acts independently of type I IFN. Additionally it was observed that the antiviral functions of Activin A does not require signalling via the type I IFN receptor (IFNAR), being unaffected by ruxolitinib (16) a pharmacological inhibitor of the JAK1 kinase downstream of IFNAR [supplementary figure 1 A-C].

**Figure 3:**
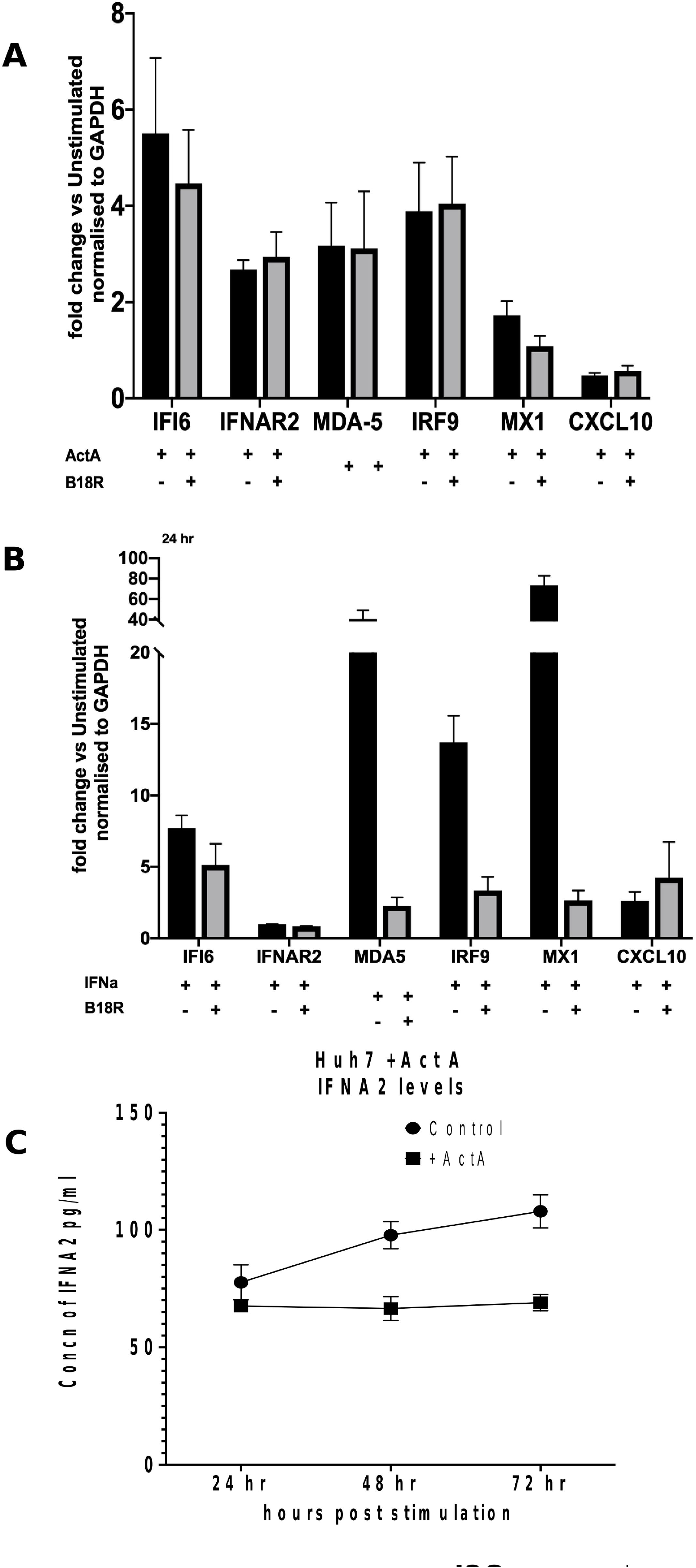
A) Gene Set Enrichment Analysis performed on two sets of genes obtained by RNA sequencing and microarray analysis. Results show high correlation in gene expression by both methods of analysis. Metacore analysis showing genes from significant canonical pathways in hepatocytes stimulated with Activin A, BMP6 and IFNa respectively. Metacore analysis showing genes from significant process network in hepatocytes stimulated with Activin A, BMP6 and IFNa respectively

### Evidence of *in vivo* expression of Activin A in viral infection

To further examine whether Activin A is expressed during the course of viral infection in vivo, we performed PCR to detect Activin A mRNA expression levels in livers and ELISA to detect Activin a protein in serum of patients with HCV infection. **Fig 4A**. shows that Activin A levels in inflamed liver samples from HCV patients were significantly increased. Significant increases in Activin A protein levels in the non-responders to IFN therapy were observed compared to controls (Fig. **4B**). To assess whether Activin A could also be induced in an acute viral infection we inoculated mice with MCMV intravenously. In this model Activin A was detected at 3 days post-infection (Fig 4C). These data indicate that acute and chronic viral infections both lead to induction of Activin A *in vivo*.

**Figure 4:**
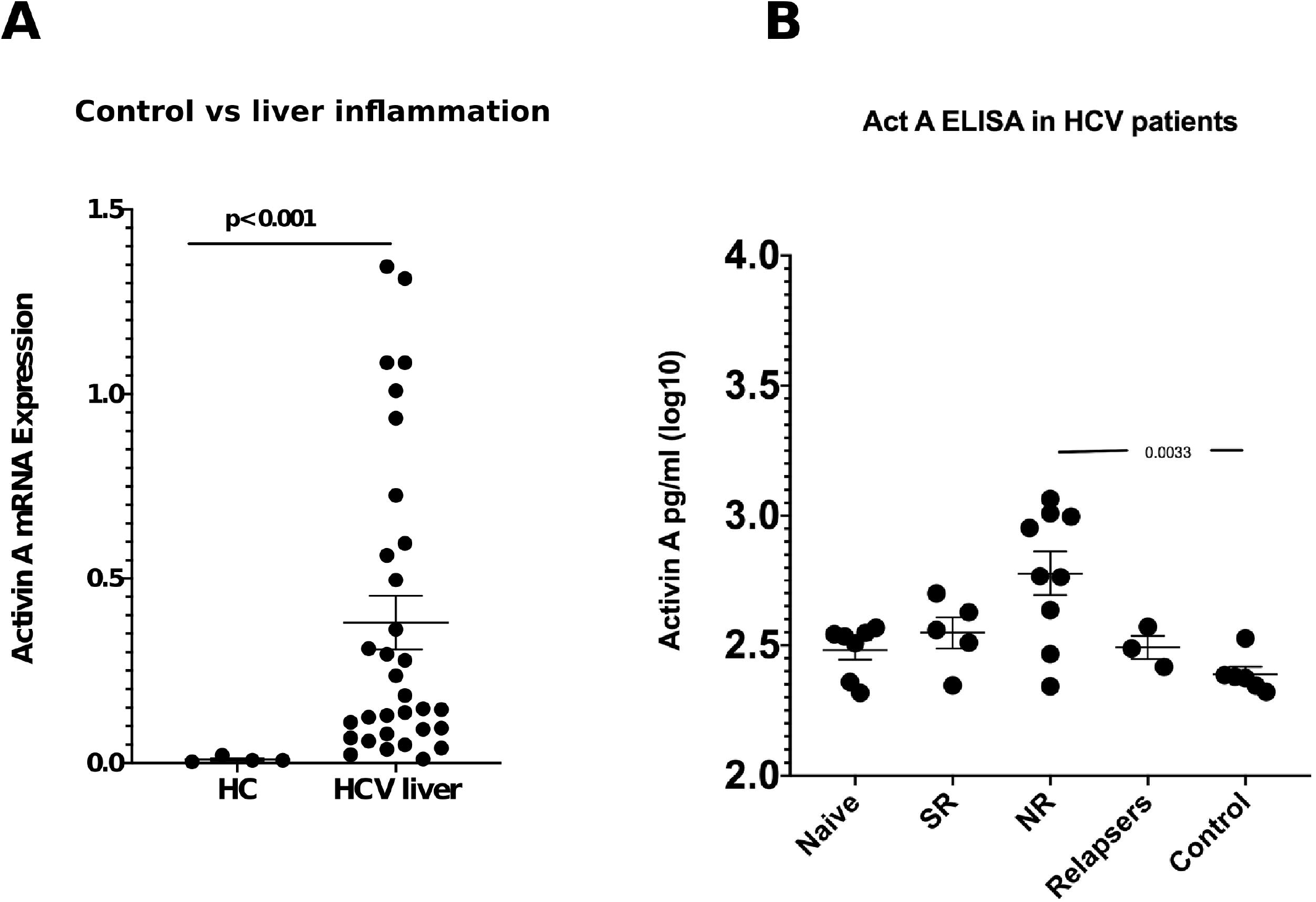
**A)** Activin A mRNA expression in patients with liver inflammation due to HCV infection, each assay done in duplicate. Patients were graded based on their ISHAK scores n=32. P-values obtained via Mann-Whitney U-test. **B)** Activin A protein in HCV serum samples as detected by ELISA. Naive (treatment naive) (n=7), SR (spontaneous resolvers)(n=5), NR (non responders)(n=9). P-values obtained via unpaired t test.

### Activin A is induced by PRR activation in human peripheral blood mononuclear cells

In order to explore the signalling mechanisms responsible for activin A induction by infectious stimuli, freshly-isolated human peripheral blood mononuclear cells, which express a broad repertoire of cytosolic and endosomal viral sensors, were transfected transfection with IVT-RNA or poly(I:C). These stimuli, detected by RIG-I and MDA5 respectively(17), elicited a statistically significant upregulation of *INHBA* mRNA (**Fig. 5A**), mirrored by significant transcriptional induction of *MX1* and *IFI6*, a pleiotropic antiviral effector (**Fig. 5B, 5C)**.

**Figure 5:**
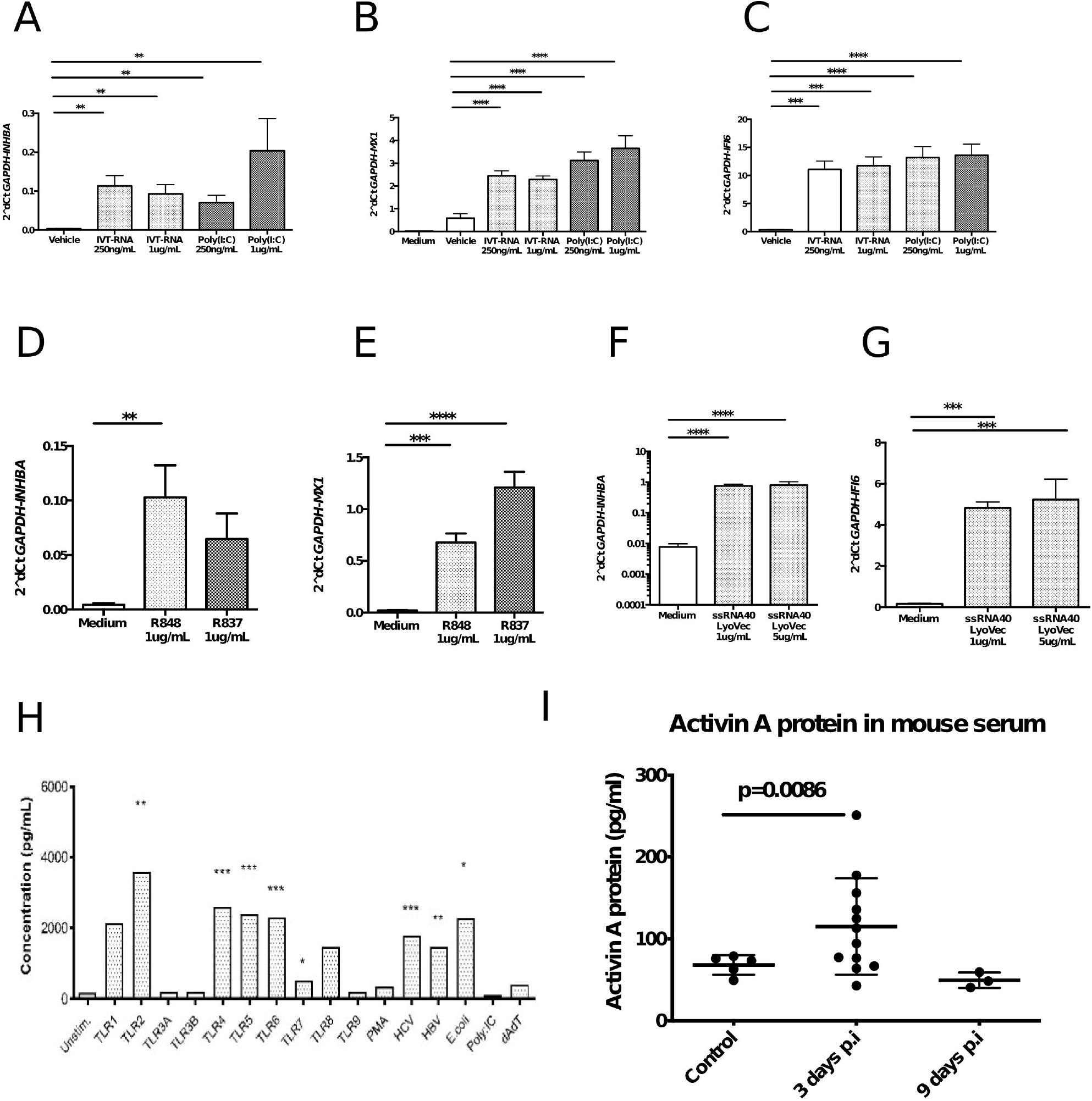
**A) B) C)** In human PBMC, *INHBA* is significantly induced by transfected poly(I:C) and poly(dA:dT) at two doses, in parallel with significant upregulation of the ISGs *MX1* and *IFI6* across all transfection conditions. **D) E)** *INHBA* is significantly induced by R848 exposure in human PBMC, with non-significant upregulation elicited by R837. *MX1* is significantly induced by both ligands. **F) G)** In human PBMC, both *INHBA and MX1* message are significantly elevated following transfection with the TLR8 agonist ssRNA40. For *INHBA* quantification, n=9 donors; for *MX1*, n=6 donors; for *IFI6*, n=3 donors. Samples transfected in singlicate; mean+SEM presented; analysis by multiple unpaired, two-tailed t-tests. **H)** Activin A protein levels as determined by ELISA on supernatants of CD14+ cells from n=3 donors stimulated with TLR agonists for 24 hrs. **I)** Activin A levels in serum of mice infected with MCMV.

Incubation with the synthetic guanidine base analogues R837 and R848 (TLR7 and TLR7/8 stimuli respectively) at 1 μg/mL resulted in approximately 15-fold and 50-fold increases in *INHBA* expression respectively **(Fig. 5D)**, implying a role for TLR7 in *INHBA* induction. This was parallel by robust induction of *MX1*, encoding the broadly antiviral GTPase MX1 (**Fig. 5E)**. Statistically significant *INHBA* upregulation following incubation with ssRNA40 at 1μg/mL and 5 μg/mL **(figure 5F)**, with attendant induction of *IFI6* (**Fig. 5G)**, suggesting a role for TLR8 in *INHBA* induction *in vitro*.

Recapitulating these observations with live viral inocula, incubation of human CD14+ monocytes with HBV and HCV (multiplicity of infection 0.5-1.0) for 16 hours resulted in significant induction of *INHBA* mRNA and detectable activin A protein in the culture supernatant (**Fig. 5H**).

### *INHBA* mRNA is upregulated by ssRNA virus infection *in vitro*

Having clarified the similarity of activin A and type I IFN in terms of patterns of antiviral gene regulation in clinical infection models and also looked at its induction by synthetic immune stimuli, our attention turned to the possibility that activin itself is induced by viral infection. In RIG-I/TLR3 replete lung adenocarcinoma-derived A549.gfp cells, infection with the ssRNA paramyxovirus Sendai virus (SeV) results in a statistically significant, dose-dependent induction of *INHBA* mRNA [figure 6A], paralleled by a strong induction of *MX1* [figure 6B]. SeV encodes a 546nt immunogenic motif within its 5’-UTR and intact RIG-I, but not MDA5, signalling is necessary for IRF3 phosphorylation and subsequent ISG induction *in vitro*(18).

**Figure 6:**
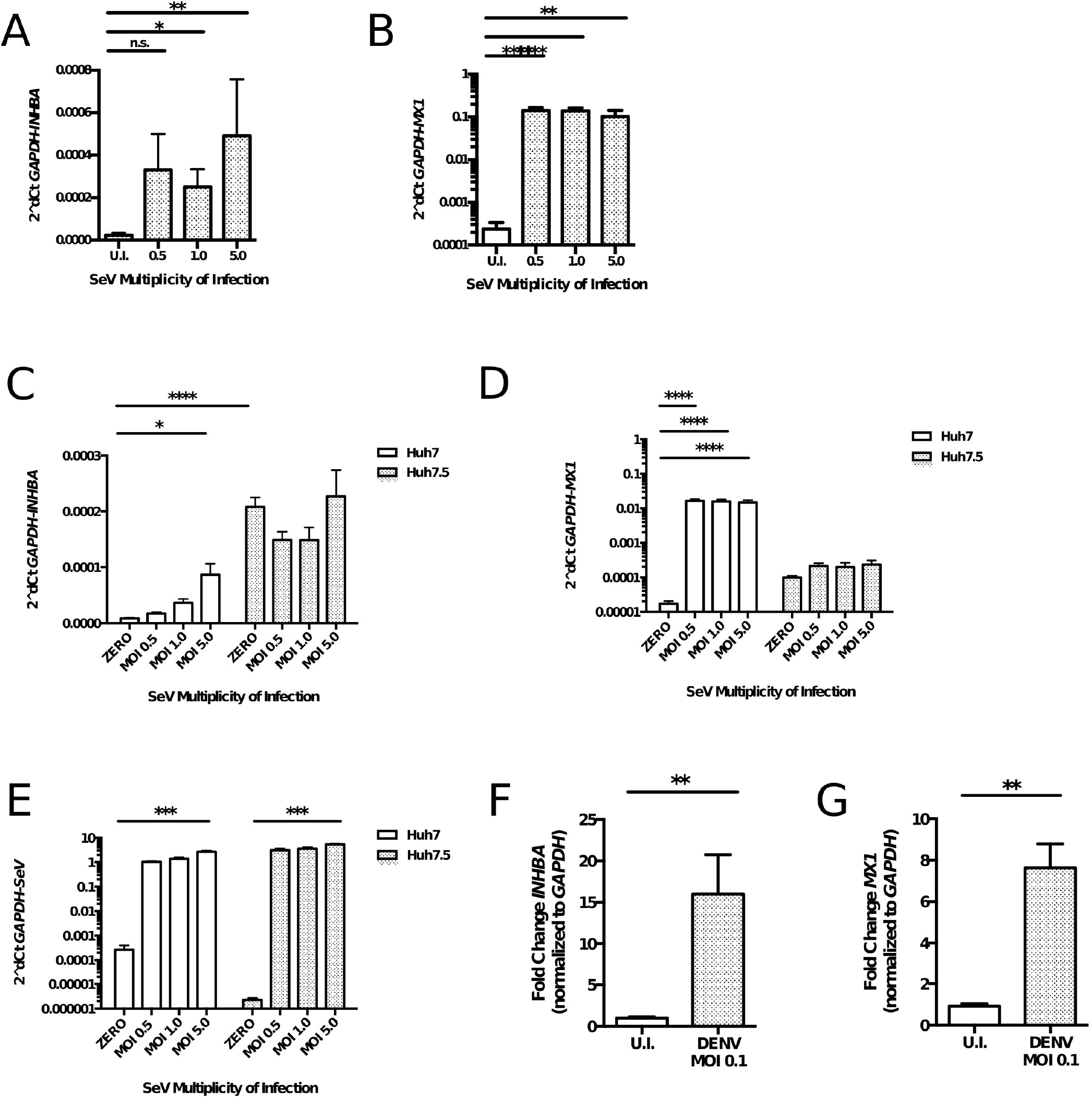
**A) B)** Both *INHBA* and *MX1* expression are significantly induced in A549.gfp cells infected with increasing titres of SeV compared to uninfected controls, at 48 h.p.i. For both *INHBA* and *MX1*, n=3 independent experiments, each in biological duplicate; mean+SEM; analysis with ordinary one-way ANOVA and Bonferroni’s post-test with respect to the uninfected condition. **C) D) E)** At 48 h, in the Huh7 line, *INHBA* mRNA is dose-dependently induced by increasing SeV titre, whereas no induction occurs in RIG-I deficient Huh7.5 cells. In parallel, *MX1* is profoundly upregulated upon SeV infection of Huh7 cells, but is not elevated in the RIG-I deficient Huh7.5.1 line (B). SeV genomic RNA is detected at robust levels upon infection at all titres in both cell line. n=2 independent experiments, each in biological duplicate; mean + SEM; analysis by two-way ANOVA and Bonferroni’s multiple comparisons test with respect to uninfected cells. **F) G)** Both *INHBA* and *MX1* are significantly upregulated in DENV-infected Huh7.5.1 cells relative to uninfected (U.I.), at 72 h.p.i. For *INHBA*, n=4; for *MX1*, n=3 independent experiments, each in biological triplicate; mean+SEM; analysis by unpaired two-tailed t-tests.

To further clarify the PRR requirements for *INHBA* induction by SeV, Huh7 and Huh7.5 hepatoma-derived cells were infected with a titration of SeV, and expression of *INHBA, MX1* and genomic SeV quantified by RT-qPCR. The RIG-I isoform expressed by Huh7.5 encodes an N-terminal point mutation that uncouples viral sensing from downstream signalling (19). Huh7 cells and their derivatives are therefore deficient for TLR3 expression.

In RIG-I replete Huh7 cells, SeV infection elicits titre-dependent *INHBA* upregulation [figure 6C], paralleled by strong induction of the IFN-dependent target gene *MX1* at 24 hours post infection [figure 6D]. In Huh7.5.1 cells, however, neither *INHBA* nor *MX1* message levels were altered by SeV infection. Baseline expression of both *INHBA* and *MX1* is approximately 10-fold higher in uninfected Huh7.5.1 cells versus Huh7. SeV genomic RNA levels were comparable in both cell lines, although the signal in the uninfected conditions implies low-level off-target amplification [figure 6E].

Moving on to consider *in vitro* flavivirus infection, type I IFN induction secondary to dengue virus (DENV) is reportedly contingent upon MAVS-dependent RLR signalling(20). MDA5 is likely to represent the principal sensor of DENV in Huh7.5.1 cells, being deficient in functional RIG-I and TLR3(18,19). At 48 hours post infection, DENV infection in Huh7.5.1 cells results in statistically significant upregulation of mRNA encoding *INHBA* [figure 6F], accompanied by a significant upregulation of *MX1* [figure 6G].

We quantified *INHBA* expression in Huh7 cells incubated with recombinant IFNα-2a, and established that *INHBA* mRNA is not induced by recombinant type I IFN in this cell line [supplementary figure 1A]. Akin to the antiviral enhancement of type I IFN by BMP6, co-incubation of OR6 HCV genomic replicon cells with recombinant activin A synergistically enhances the antiviral effect of IFNα-2a, hinting at possible functional ramifications of viral *INHBA* induction [supplementary figure 1 B-D].

### Influenza A infection induces *Inhba* expression in the lungs of C57BL/6 mice

Moving to *in vivo* infection models, we observed that *Inhba* expression is significantly upregulated in the lungs of C57BL/6 mice infected with the influenza A [FLUAV] strain PR/8/34 for 72 hours compared to uninfected controls [figure 7A]. This induction correlates tightly with that of *Isg15*, encoding a pleiotropic, IFN-stimulated antiviral effector(21) (R^2^= 0.9271, p=0.0020) [figure 7B-7C]. In the liver, however, *Inhba* expression is significantly suppressed by FLUAV [figure 7D], despite no alteration in *Isg15* expression [figure 7E]. Suppression of hepatic *Inhba* expression in the presence of a systemic inflammatory response has been previously reported, albeit in the context of LPS administration (22,23), not viral infection.

**Figure 7:**
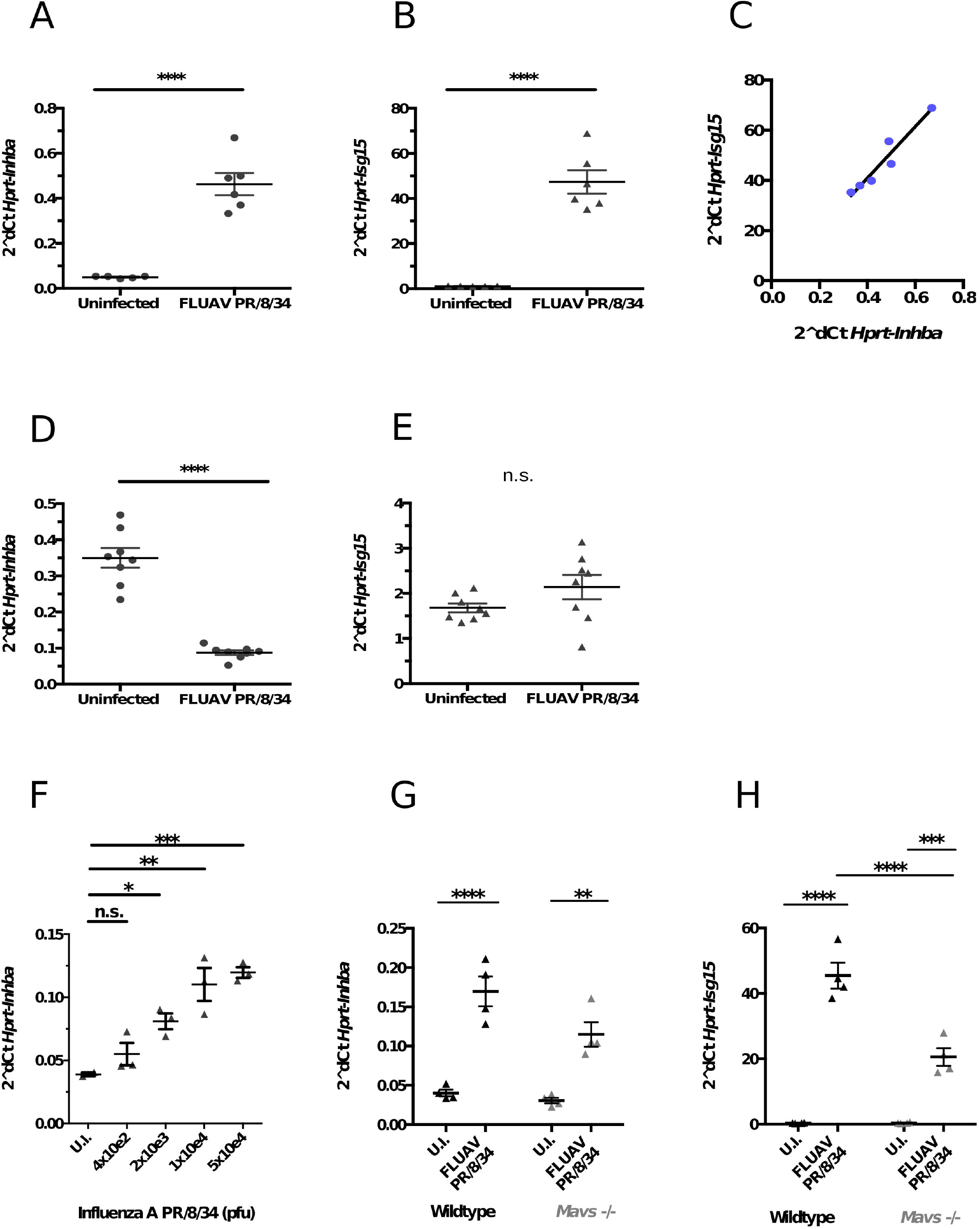
**A) B) C)** *Inhba* and *Isg15* mRNA are upregulated in the lungs of FLUAV PR/8 infected mice at 72 h.p.i. The degree of their respective induction is strongly positively correlated (R2= 0.9271, p=0.0020). n=5 uninfected and n=6 infected mice; mean+SEM; analysis by unpaired two-tailed t-test (A,B) and linear regression (C). **D) E)** At 72 h.p.i. in the liver, *Inhba* message is significantly suppressed in the infected mice, whereas hepatic levels of *Isg15* message do not differ between groups. n=8 mice per group; mean+SEM; analysis by unpaired two-tailed t-test. **F)** Lung *Inhba* expression is dose-dependently induced by FLUAV infection at 48 h.p.i. n=3 mice per group; mean+SEM; analysis by ordinary one-way ANOVA and Bonferroni’s post-test with respect to uninfected mice **G) H)** mRNA encoding both *Inhba* and *Isg15* is significantly upregulated in the lungs of both WT and Mavs KO mice at 48 h.p.i. with FLUAV; no significant difference in the magnitude of induction is evident between groups.

Upon intranasal infection with titrating doses of FLUAV, *Inhba* is dose-dependently upregulated at 48 hours post infection, reaching maximal a three-fold increase with the 5×10^4^ plaque forming unit (pfu) inoculum [figure 7F].

To gain a mechanistic insight into the PRR subsets responsible for *Inhba* induction during intranasal FLUAV infection, we compared *Inhba* expression in lung lysates from both wildtype (WT) and Mavs knockout (KO) mice. These animals cannot transduce RLR signalling (24). In wildtype and Mavs KO mice, *Inhba* expression was significantly increased by intranasal administration of 5×10^4^ pfu FLUAV at 48 hours post-infection,albeit by a smaller fold change in the transgenic animals [figure 7G]. Similarly, the magnitude of both *Inhba* and *Isg15* induction was lower in Mavs KO compared to WT mice, although this difference did not attain statistical significance [figure 7H]. These data suggest a function of Mavs-independent pattern recognition in *Inhba* induction by FLUAV.

## Discussion

An antiviral function of the BMP/activin-SMAD signalling axis has been described in the context of viral infection(6). Here, we show that gene expression signatures associated with activin stimulation are induced in parallel to those downstream of type I IFN in a clinically important viral disease, namely chronic HCV infection. Additionally, we provide evidence for induction of activin A, encoded by the *INHBA* gene, in multiple models of viral infection, both *in vitro* and *in vivo*.

The transcriptomic analyses described here suggest that stimulation of antiviral effector genes classically regarded as “interferon-stimulated” may not be an exclusive feature of IFN, as both BMP6 (6) and activin A are able to upregulate ISG in the absence of exogenous IFNα and ISG induction occurs as a summation of diverse unrelated signalling events. Together with the observed antiviral properties of recombinant activin A, we propose that its transcriptional induction downstream of viral sensing may represent a hitherto undescribed antiviral feedback response, initiated upon detection of infection and acting to curtail viral replication.

Wu and colleagues report that in the resting state, murine alveolar and bronchial macrophages stain positively for activin A protein; during LPS endotoxaemia, activin A positive neutrophils aggregate in the perialveolar spaces (23). This reported accumulation of activin A-rich neutrophils in the lung correlated with reduced numbers in the bone marrow, consistent with migration into the systemic circulation. It was posited that this activin A induction occurred at the post-transcriptional level, being ablated by cyclohexamide but unaffected by the transcription initiation inhibitor actinomycin D (23). In contrast, our data obtained with FLUAV demonstrate transcriptional upregulation of *Inhba*, with respiratory epithelia and alveolar macrophages being equally plausible candidate sources.

*In vivo, Inhba* mRNA is upregulated in the lungs of mice following intranasal FLUAV infection. This induction is not abrogated upon Mavs knockout, implying a contribution of Mavs-independent viral sensing. Data from human PBMC *in* vitro indicate that TLR7/8 activation – that is, independent of MAVS – is sufficient for *INHBA* upregulation. As such, definitive characterization of *Inhba* induction *in vivo* will require examination of mice deficient in Tbk1, the point of RLR and TLR convergence upon IRF3.

Our observations of *INHBA* induction in experimental infection systems correspond with clinical reports of elevated serum activin A protein in both acute influenza A infection, where it correlates with the degree of respiratory distress, and in chronic HCV (5,25). Interpretation of these patient studies is complicated by the well-established roles of activin A and TGFβ signalling in tissue fibrosis (26); it is impossible to dissect whether elevated activin A occurs immediately downstream of viral sensing or as a mediator of fibrotic immune pathology.

Given the described antiviral properties of exogenous activin proteins and manipulation of the activin/SMAD signalling axis(6), these data nevertheless provide an essential first step in the clarification of activin A’s endogenous role in the innate response to viral infection.

## Supporting information

Table 1: Top 40 Differentially-Regulated Genes

Supplementary Table 1: PCR Primer Sequences

## Acknowledgements

The authors would like to thank Dr. Azim Ansari for useful discussions; Dr Jo Miller, Professor Nicole Zitzmann, Dr Rachel Rigby and Professor Jan Rehwinkel for technical advice and experimental assistance. The authors also thank the High-Throughput Genomics Group at the Wellcome Trust Centre for Human Genetics (funded by Wellcome Trust grant reference 090532/Z/09/Z and MRC Hub grant G0900747 91070) for the generation of the Sequencing and micro-array data). This research was supported by the Wellcome Trust (WT109965MA) NIHR Senior Fellowship (PK), NIH U19 I082360. The authors wish to acknowledge the BRC Oxford GI Biobank in collecting and making available the samples/data used in the generation of this publication. The research was supported by the National Institute for Health Research (NIHR) Oxford Biomedical Research Centre (BRC). The views expressed are those of the authors and not necessarily those of the NHS, the NIHR or the Department of Health.

The data that support the findings of this study are available from the corresponding author upon reasonable request.

## Author contributions

KAH, NR, PK and HD designed the experiments. KAH, NR and RME performed the experiments and wrote the manuscript, LNL performed *in vivo* MCMV infections, PF provided liver biopsy tissue and EM performed the bioinformatic analysis.

## Materials and Methods

### Transcriptomics pipeline: extraction and sample preparation

Total RNA was prepared after stimulation or that of untreated controls with the RNeasy Mini/ Plus Micro Kit (Qiagen, UK cat# 74106/ 74034). Stimulated cells were lysed in buffer RLT (Qiagen) and homogenized with a QIAshredder (Qiagen). RNA was quantified spectro-photometrically and the quality of RNA preparations for Illumina microarray analysis was checked with an Agilent Technologies 2100 Bio-analyzer.

Two-step reverse transcription was performed on the RNA using AppScript cDNA synthesis kit (Appleton Woods) and quantitative PCR performed using the Roche Light Cycler 480. Approximately 500ng of total RNA from cells was reverse transcribed using the AppScript cDNA synthesis kit. Briefly, total RNA was mixed with kit components App RTase and App cDNA mix to a final volume of 20microlitrel. The mix was then incubated at 42°C for 30 mins. At the end of the reaction RTase was inactivated by heating the reaction to 85°C for 10 mins.

For RT-qPCR validation, 5ul of a 1/20th dilution of resultant cDNA was used as a template for qPCR using the Roche light cycler 480 to detect expression of selected ISGs. Primers were designed using the Roche Universal Probe Library system. Relative gene expression was calculated using the comparative cycle threshold method (27) normalized to expression of the house-keeping gene GAPDH and expressed relative to a mock treated sample.

### Transcriptomics pipeline: microarray analysis

500ng of total RNA was used for microarray analysis and was performed by the Oxford Genomic Centre at the Wellcome Trust Centre for Human Genetics, University of Oxford. The quality of the RNA was to have a RNA integrity number (RIN) greater than 7 and 28S/18S ratio of greater than 1.6.

A whole genome gene expression analysis was performed utilizing the Human HT12v4.0 Expression Beadchip. The RNA was converted to biotin labelled cRNA which was then hybridised to the chip. The hybridised chip was then scanned using llumina’s iScan scanner. The gene expression profile was then created using illumina’s GenomeStudio software.

### Transcriptomics pipeline: RNA sequencing analysis

The mRNA fraction was selected from the total RNA provided before conversion to cDNA. dUTP was incorporated during second Strand cDNA synthesis. The cDNA was end-repaired, A-tailed and adapter-ligated. Before amplification the samples were uridine digested. The prepared libraries were then size selected, multiplexed and QC’ed before paired end sequencing over one rapid run. The data was then aligned to the reference and quality checked.

### Cell lines

OR6 cells (kindly from Prof Raymond Chung [MGH, MA, USA] with permission of Prof Kato and Dr Ikeda [Okayama University, JP]) were cultured in Dulbecco’s Modified Eagle’s Medium (DMEM), supplemented with 10% Foetal Calf Serum (FCS) (PAA,, AT or Lonza, CH), 100U/mL penicillin (Sigma-Alrdich, USA), and 0.1mg/mL streptomycin (Sigma-Aldrich) and 2mM L-glutamine (Sigma-Aldrich): hereafter referred to as “D10”. OR6 cells are stably transfected with a full-length HCV genomic replicon in tandem with a Renilla luciferase reporter and neomycin resistance cassette. OR6 cells were under negative selection with G418 100 μg/mL (Sigma-Aldrich).

Huh7 (ATCC, USA) and Huh7.5.1 (kindly from Prof Raymond Chung [MGH, MA, USA]) cells were maintained in D10.

HepaRG were cultured at 37°C, 5% CO_2_ in Williams’ E medium (with Glutamine) (Gibco-Invitrogen) supplemented with glutamine (2 mM), penicillin/streptomycin (50 U/mL), gentamycin (20 µg/mL), insulin bovine (5 µg/mL, Roche-Boehringer-Manheim, France), hydrocortisone hemisuccinate (7×10^−5^ M, Roche-Boehringer-Manheim) and FCS (10% selected, non-decomplemented, Fetaclone II-Hyclone-PERBIO France). The cells were allowed to differentiate in the presence of 2% (v/v) DMSO to the medium.

A549.gfp cells (a kind gift from Prof Richard Randall, University of St Andrews, UK) were maintained in in D10.

### Patients Samples

HCV patient samples were obtained from patients enrolled at the JR Hospital, Oxford and samples stored under protocol 16/YH/0247. Liver biopsy samples were collected prior to commencement of anti-viral therapy and were graded and staged using ISHAK scoring at S. Bortolo Hospital Vicenza, Italy. RNA extraction, reverse transcription and qRT-PCR was performed as described below. Ethical approval for use of the biopsy samples was obtained from the relevant local ethics committees. Informed written consent was obtained from all patients involved.

### Peripheral blood mononuclear cells extraction

Whole blood was procured from nine healthy volunteer donors in accordance with local ethical policy and practices, as described previously(28,29). Blood samples were heparin-treated before immediate layering onto a Ficoll-Pacque Plus™ (GE Healthcare Life Sciences, USA) density centrifugation gradient in a 1:1 volume ratio. Samples were centrifugedand the buffy coat layer aspirated with a pipette. Buffy coat cells were washed three times in 2.5 mM EDTA-PBS at 4°C before dilution and maintenance in RPMI-1640 supplemented with 10% FCS (PAA, AT or Lonza, CH) 2 mM glutamine, 100 U/mL penicillin, 0.1 mg/mL streptomycin (all Sigma-Aldrich).

### Enrichment of CD14 monocytes

CD14 positive cells were isolated from the PBMCs using the EasySep™ Human CD14 Positive Selection Kit by STEMCELL Technologies™ (Cat# 18058).

### Synthetic nucleic acids and base analogues

High molecular weight poly(I:C) [cat I.D. tlrl-pic], poly(dA:dT) [cat. I.D: tlrl-patn], *E. coli* K12 dsDNA [cat. I.D. tlrl-ecdna], ODN 2216 [cat. I.D. tlrl-2216], R837 [cat. I.D. tlrl-imqs], R848 [cat. I.D. tlrl-r848] and ssRNA40/LyoVec™ [cat. I.D. tlrl-lrna40] were purchased from InvivoGen (FR). IVT-RNA transcribed by the T7 RNA polymerase and corresponding to nucleotides 1-99 of the neomycin phosphotransferase gene was synthesized as described in (30) (kindly from Prof Jan Rehwinkel [University of Oxford, UK]).

### Transfection and stimulation of unfractionated PBMC and CD14+ monocytes

Human PBMC were extracted from healthy donors as described in section 2.1.6 and plated at 1×10^6^ cells/well of a 24-well plate. PBMC were transfected with IVT-RNA and poly(I:C) using Lipofectamine® LTX with PLUS™ (Life Technologies). R837, R848, ODN 2216 and ssRNA40 were added to the culture medium in the absence of transfection reagent to the end-point concentrations specified.

CD14+ cells were plated at at 1×10^6^ in 300 µl R10 medium per well and stimulated with HBV inoculum (final MOI = 0.5-1), HCV inoculum (final MOI = 0.5-1). Cells were incubated at 37°C, 5% CO_2_ for 16 hours. Cells and supernatants were harvested and frozen at -20°C for further analysis.

### *In vitro* Sendai virus (SeV) Infection

Cantell strain Sendai Virus [cat. I.D. ATCC® VR-907™] was purchased from ATCC. A549, Huh7 or Huh7.5 cells were plated at 5×10^4^ cells/well in 12-well plates and incubated at 37°C for 24h. Immediately prior to infection, D10 was aspirated and the monolayers washed twice times with PBS, before addition of SeV at MOI 0.5, MOI 1 and MOI 5 in 500uL serum-free DMEM. Cells were incubated with SeV for 2h at 37 °C, before removal of the virus, two PBS wash/aspirations and replenishment with D10.

### *In vitro* dengue virus (DENV) infection

Huh7.5 cells were plated at 3×10^5^ cells/well in 12-well plates and allowed to adhere. Cells were incubated with DENV2 (strain 16681) in serum-free DMEM for 2 h at room temperature with gentle agitation, followed by removal of virus, washing with PBS and incubation at 37 °C for 48h in D10.

### *In vivo* influenza A virus (FLUAV) infection

For figure 4A-E, 6 week old female C57BL/6 mice were anaesthetized and infected intranasally (i.n.) with 3.5 haemagglutinating units FLUAV PR/8/34 (H1N1), kindly provided by Prof John Skehel (NIMR, Mill Hill, UK). Control mice were administered PBS i.n. Mice were sacrificed at 72 h.p.i. by asphyxiation. Whole lungs were immediately lysed in RLT buffer (Qiagen) with 10 μL/mL β-mercaptoethanol and mechanically disrupted with a TissueRuptor (Qiagen), before RNA extraction, cDNA synthesis and RT-qPCR quantification of gene expression as described in section 2.2. Liver explants (∼2mm^3^) were preserved in RNAlater (Qiagen), before mechanical lysis with a TissueRuptor and RNA extraction, cDNA synthesis and RT-qPCR quantification of gene expression.

For figure 4F-G, 6-week old female wildtype C57BL/6 mice were infected i.n. with increasing titres of wildtype FLUAV PR/8/34 from 4×10^2^ to 5×10^4^ pfu. Control mice were administered DMEM i.n. Mice were sacrificed at 48 h.p.i. by asphyxiation, followed by RNA extraction, cDNA synthesis and RT-qPCR quantification of gene expression. After sacrifice, whole lungs were removed and snap-frozen in liquid N_2_. 1 mL Tri Reagent (Sigma-Aldrich) was added to each lung, before lysis with glass beads [cat. I.D. G8772] (Sigma-Aldrich) in a FastPrep Cell Disrupter. RNA was extracted from the lungs with lysates with phenol-chloroform followed by isopropanol precipitation Sacchi (31). RNA isolates were further purified, and depleted for genomic DNA, by secondary extraction using the RNeasy Plus kit (Qiagen). cDNA synthesis and RT-qPCR analysis of gene expression were conducted as described previously.

For figure 4H-I, 6-week old female C57BL/6 mice, both wildtype and MAVS KO (32) were infected i.n. with 5×10^4^ pfu wildtype FLUAV PR/8/34. Mice were sacrificed at 48 h.p.i. by asphyxiation. Control mice were administered DMEM i.n. Whole lungs were harvested and RNA extracted as per the preceding instructions, followed by cDNA synthesis and RT-qPCR quantification of gene expression. Animal experimentation was in line with requirements stipulated in the ARRIVE guideline checklist.

### RNA isolation and quantification

Unless otherwise specified, cell lysates were homogenized with a QIAshredder column (Qiagen) and RNA extracted with the RNEasy Mini Kit (Qiagen). RNA concentration was determined with a NanoDrop 2000 Spectrophotometer (Thermo Fisher Scientific, MA, USA) at 260 nm. The quality of RNA preparations for Illumina microarray analysis and RNA sequencing analysis was checked with an Agilent Technologies 2100 Bio Analyser.

### cDNA synthesis and RT-qPCR analysis

cDNA was reverse-transcribed from template RNA either using a two-step reverse transcription using AppScript cDNA synthesis kit (Appleton Woods) or using the High Capacity RNA-to-cDNA kit (Applied Biosystems, USA).

All RT-qPCR reactions were either performed using an Applied Biosystems 7500 Fast Real-Time PCR System (Applied Biosystems, USA). For TaqMan™ quantification, gene expression was assessed with inventoried TaqMan™ Gene Expression Assays (Applied Biosystems, MA, USA) diluted in TaqMan™ Gene Expression Master Mix (Applied Biosystems, USA). TaqMan™ inventoried assays used: *GAPDH* (glyceraldehyde 3-phosphate dehydrogenase) Hs99999905_m1; *INHBA* (inhibin beta A) Hs01081598_m1; *MX1* (myxovirus resistance 1, mouse, homolog of) Hs00895598_m1; *IFI6* (interferon-alpha-inducible-protein 6) Hs00242571_m1; *IFNAR2* (Interferon alpha, beta and omega, receptor 2) Hs01022061_m1; *JAK1* (Janus kinase 1) Hs01026983_m1; *STAT2* (Signal transducer and activator of transcription 2), Hs01013123_m1; Hprt (hypoxanthine-guanosine phosphoribosyltransferase) Mm01545399_m1; *Inhba* (inhibin beta A) Mm00434339_m1; *Isg15* (ubiquitin-like modifier ISG15) Mm01705338_s1; *Trim14* (tripartite motif-containing protein 14) Mm01352552_m1 or quantitative realtime PCR analysis was performed using the Roche Light Cycler 480 instrument using AppProbe reagents (Appleton Woods). Primers were designed using the Roche Universal Probe library system. Relative gene expression was calculated using the comparative cycle threshold method(27)normalised to expression of the housekeeping gene GAPDH and expressed relative to a mock treated sample. The primer list used in fig 1 D and E, 2 A and B, 4 A are indicated in Supplementary Table 2.

### Plasma cytokine and Activin A quantification

Concentrations of interferon alpha 2 in cell culture supernatants was measured using custom multiplex immunoassay kits (GeniePlex, Ireland).

Concentration of Activin A in human and mouse serum samples were performed using sandwich ELISA-Activin A immunoassay kit from R&D system (Cat DAC008).

### Bioinformatic and Statistical Analysis

The Illumina bead chip output files were processed and analysed using the R statistical software (v 2.11)(33) and statistical testing was performed using the Linear Models for Microarray Analysis (limma) package and DESeq2(13,34). Differential gene expression between the experimental groups was assessed by generating relevant contrasts corresponding to the possible cell type comparisons. Raw *p*-values were corrected for multiple testing using the false discovery rate controlling procedure of Benjamini and Hochberg (35); adjusted p-values <0.01 were considered significant. Gene set enrichment analysis methods (GSEA) are as described(36,37). Pathway representation was performed using the MetaCore pathway analysis software from Clarivate Analytics. Unless otherwise specified, data transformation and analysis was performed with Microsoft Excel (Microsoft Inc., USA). Statistical analysis and data presentation were performed with GraphPad Prism (GraphPad Software Inc., USA).

---

**Supplementary Figure 1:**

**A)** HuH7 cells were incubated with recombinant IFNα 2a for 24h, followed by RNA extraction and RT-qPCR quantification of mRNA encoding *INHBA*. n=4 independent experiments; mean + S.E.M.; analysis by two-tailed unpaired t-test.

**B) C) D)** Co-incubation with activin A synergistically enhances the antiviral effect of IFNα in the OR6 HCV replicon model. OR6 cells were incubated with a titration of recombinant human IFNα in the presence of titrating doses of recombinant human activin A; cells were assayed at 72 h post-incubation for Renilla luciferase (A) and Cell Titer Glo (B), with signal presented relative to the untreated condition. Renilla luciferase signal is normalized to Cell Titer Glo intensity, to account for variation in cell number, in (C). Normalized for the dose-dependent diminution in cell number/viability caused by activin A, co-incubation with activin A synergistically enhances the antiviral effect of both 10 U/mL and 100 U/mL IFNα. n=3 independent experiments in biological triplicate; mean+SEM; analysis by 2-way ANOVA and Bonferroni’s multiple comparisons test with respect to the ZERO activin A condition within each group.

