## Supplementary figures and images for "Innate triggering and antiviral effector functions of activin A"

### Supplementary Figure 1

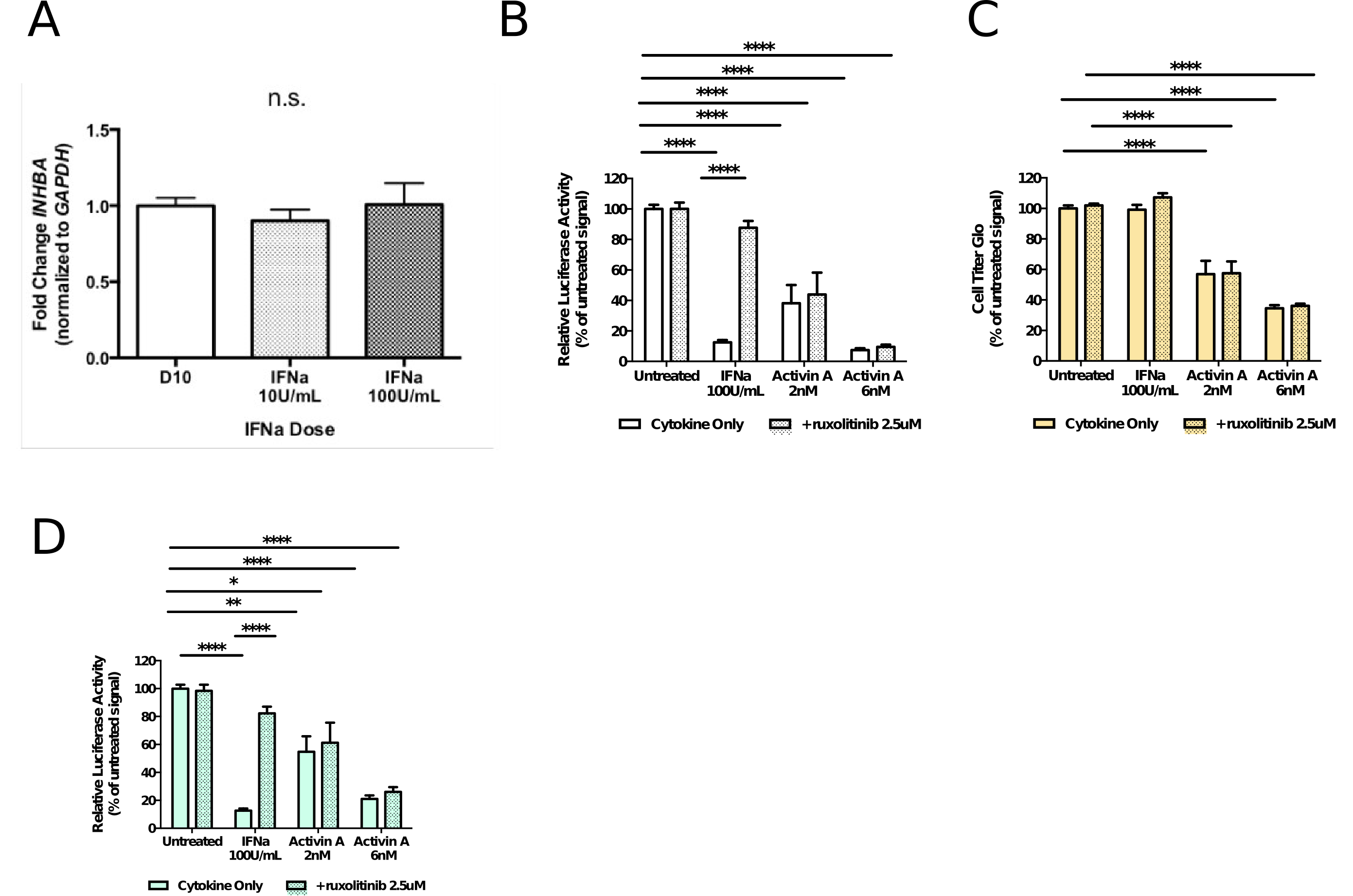

### Supplementary Table 1: PCR Primer Sequences

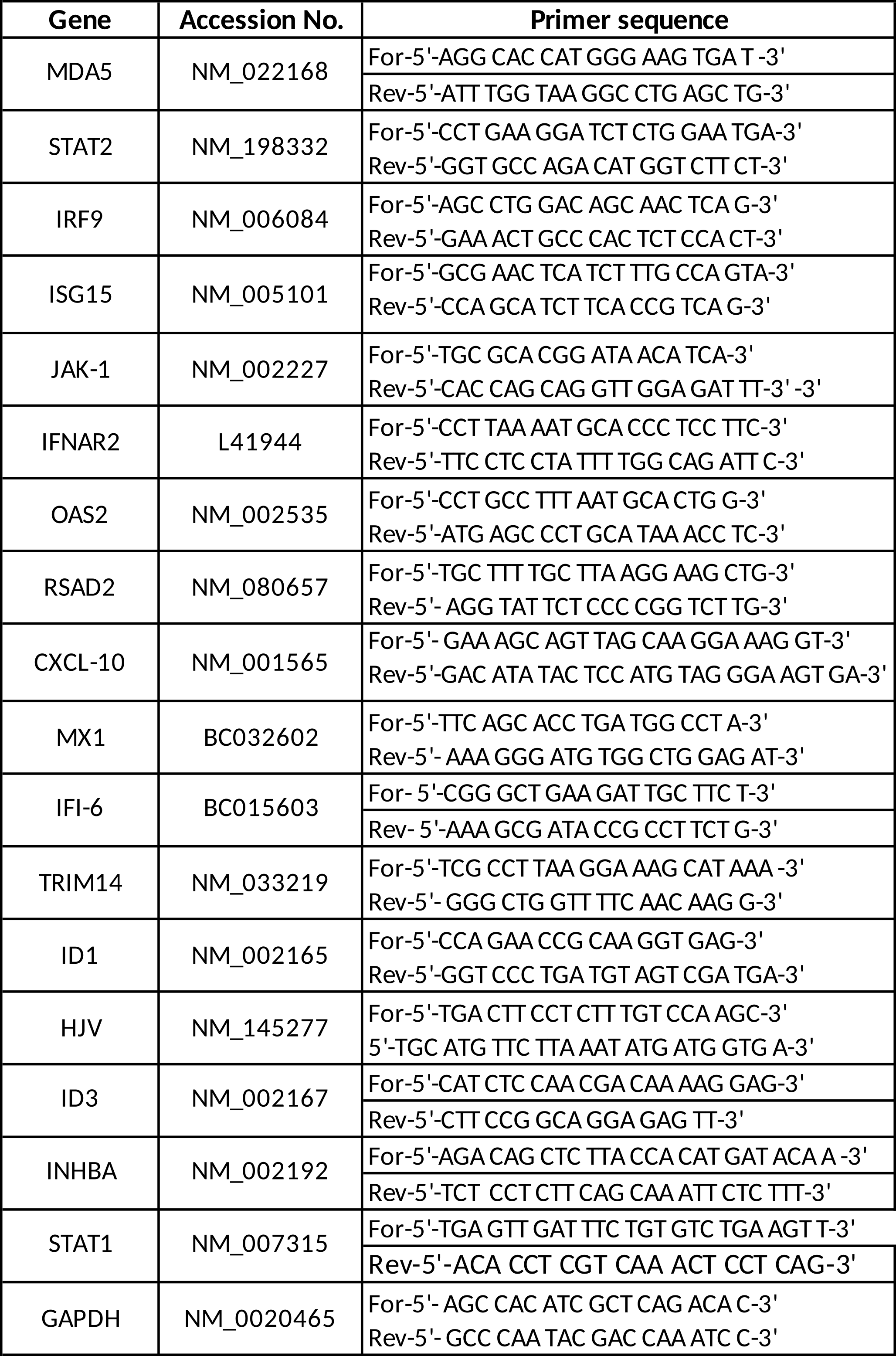

### Table 1: Top 40 Differentially-Regulated Genes

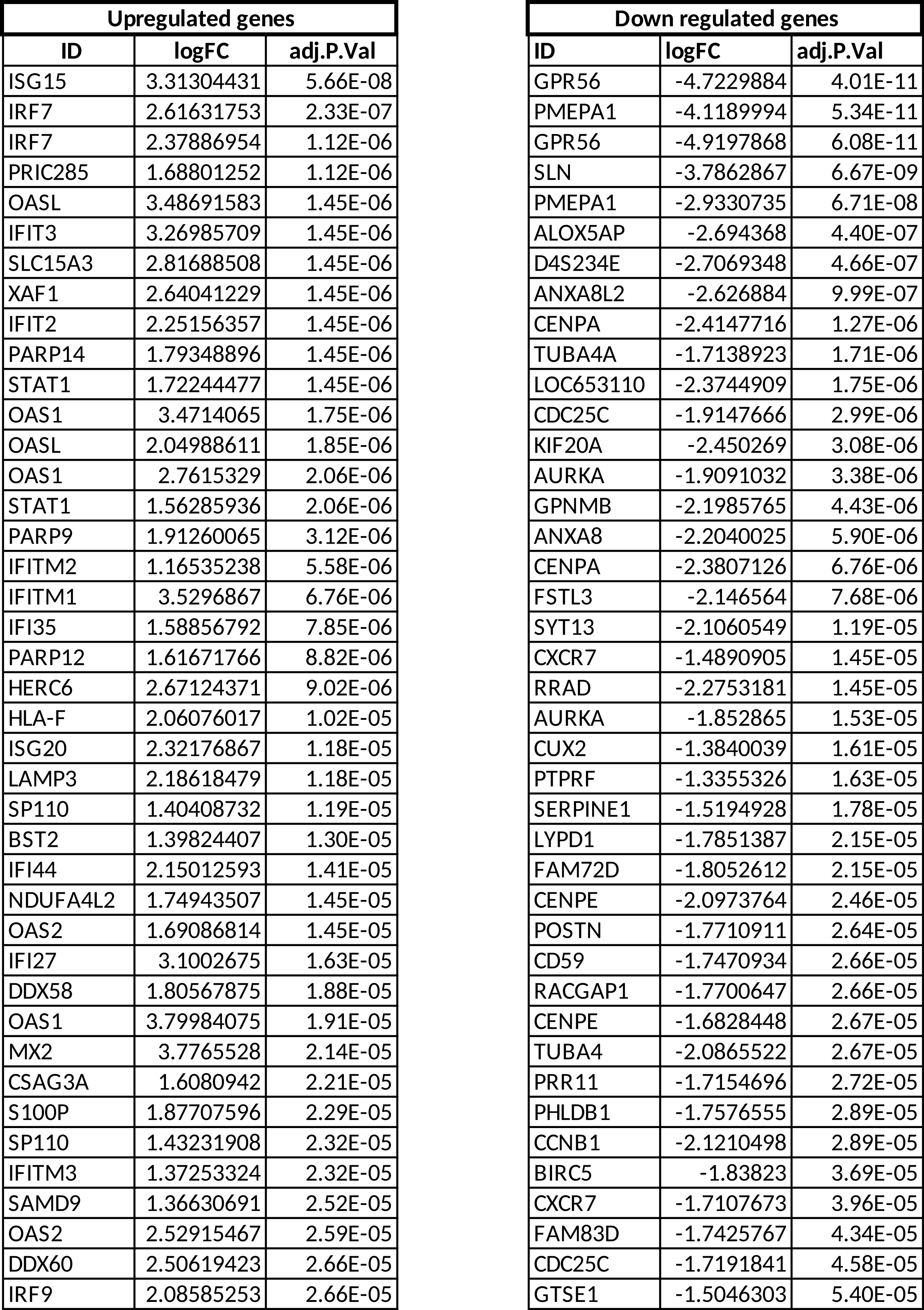
