## Supplementary Information for "Innate triggering and antiviral effector functions of activin A"

### **Supplementary materials and methods:**

**OR6 luciferase assay**

In supplementary figure 1A, OR6 cells were plated at 3.5x10^3^ cells/well in flat-bottomed 96-well plates. After 24h incubation, cells were incubated in triplicate wells with D10 (untreated), recombinant human IFNα 100U/mL (PBL Assay Science), recombinant human BMP6 (R&D) or a titration of recombinant human activin A or activin B (both R&D) from 0.25 nM to 4 nM, for 72 h. Ruxolitinib (INCB018424) is a small-molecule inhibitor of the JAK1 intracellular kinase required for signal transduction by the IFNAR2 type I IFN receptor chain(16). In supplementary figure 1B, OR6 cells were plated at 3.5x10^3^ cells/well in flat-bottomed 96-well plates, triplicate wells per experimental condition. Cells were incubated with 5 μM ruxolitinib (Tocris Bioscience UK) or D10 (untreated) in 50 μL for 30 min, followed by the addition of a unit volume recombinant human IFNα to 100 U/mL (PBL Assay Science) or recombinant human activin A (R&D Systems) to 2 nM or 6 nM. Cells were incubated for 72 h and assayed at end-point for Renilla luciferase activity. End-point measurement of luciferase activity was performed with the Renilla-Glo Luciferase Assay System (Promega, USA). Measurements were obtained with a Fusion II Luminometer (PerkinElmer, USA) or a GloMax 96 Microplate Luminometer (Promega, USA).

**Cell Titer Glo © Assay**

Cells were assayed for intracellular ATP content, an index of cell number and viability, with the Cell Titer Glo Assay System (Promega, USA). Lysates were extracted as per the kit’s protocol and transferred to an opaque plate before measurement of luminescence intensity.
